# Massive transient damage of the olfactory epithelium associated with infection of sustentacular cells by SARS-CoV-2 in golden Syrian hamsters

**DOI:** 10.1101/2020.06.16.151704

**Authors:** Bertrand Bryche, Audrey St Albin, Severine Murri, Sandra Lacôte, Coralie Pulido, Meriadeg Ar Gouilh, Sandrine Lesellier, Alexandre Servat, Marine Wasniewski, Evelyne Picard-Meyer, Elodie Monchatre-Leroy, Romain Volmer, Olivier Rampin, Ronan Le Goffic, Philippe Marianneau, Nicolas Meunier

**Author notes:** Corresponding author Nicolas Meunier (VIM, INRA-Biotechnologies Bat 440, 78352 Jouy-en-Josas Cédex).

## Abstract

Anosmia is one of the most prevalent symptoms of SARS-CoV-2 infection during the COVID-19 pandemic. However, the cellular mechanism behind the sudden loss of smell has not yet been investigated. The initial step of odour detection takes place in the pseudostratified olfactory epithelium (OE) mainly composed of olfactory sensory neurons surrounded by supporting cells known as sustentacular cells. The olfactory neurons project their axons to the olfactory bulb in the central nervous system offering a potential pathway for pathogens to enter the central nervous system by bypassing the blood brain barrier. In the present study, we explored the impact of SARS-COV-2 infection on the olfactory system in golden Syrian hamsters. We observed massive damage of the OE as early as 2 days post nasal instillation of SARS-CoV-2, resulting in a major loss of cilia necessary for odour detection. These damages were associated with infection of a large proportion of sustentacular cells but not of olfactory neurons, and we did not detect any presence of the virus in the olfactory bulbs. We observed massive infiltration of immune cells in the OE and *lamina propria* of infected animals, which may contribute to the desquamation of the OE. The OE was partially restored 14 days post infection. Anosmia observed in COVID-19 patient is therefore likely to be linked to a massive and fast desquamation of the OE following sustentacular cells infection with SARS-CoV-2 and subsequent recruitment of immune cells in the OE and *lamina propria*.

## 1. Introduction

Respiratory viruses are very common worldwide and impact human and animal health with important economic consequences. Although most symptoms of respiratory virus infection are related to airway inflammation and focused on the respiratory tract, extra respiratory organs are also targeted by these viruses including the central nervous system (Bohmwald et al., 2018). The recent pandemic of coronavirus disease emerging in December 2019 (COVID-19) has been accompanied by a prevalence of olfaction alteration. Indeed, approximately 60% of patients infected by the severe acute respiratory syndrome coronavirus 2 (SARS-CoV-2) that is the cause of COVID-19 suffered from mild (hyposmia) to total loss of olfactory capacities (anosmia) (Spinato et al., 2020). The origin of these alterations remains to be established.

Olfaction starts with olfactory sensory neurons (OSN) located in the olfactory epithelium (OE) present in the most dorsal part of the nasal cavity. The OE along with the underlying *lamina propria* constitutes the olfactory mucosa. OSNs have cilia in direct contact with the environment in order to detect odorants. They are therefore accessible to respiratory viruses and some of these viruses are able to infect the OSN and reach the central nervous system by following the olfactory nerves which project to the olfactory bulbs (Forrester et al., 2018; Bryche et al., 2019a). Interestingly, SARS-CoV-1 (Netland et al., 2008) was described as a virus able to use this olfactory pathway.

ACE2 (angiotensin-converting enzyme 2) was characterized as the main entrance receptor for SARS-CoV-2 (Letko et al., 2020), similarly to SARS-CoV-1. SARS-CoV-1 was described as being able to enter the central nervous system in mice transgenic for human ACE2 (Netland et al., 2008). In this model, the brain infection starts from the olfactory bulbs and while the authors did not look for a potential infection of the olfactory mucosa, it may be that SARS-CoV-1 could infect olfactory sensory neurons, reaching the brain through the olfactory bulbs where these neurons project (Forrester et al., 2018). If SARS-COV-2 infection follows the same pathway, anosmia following SARS-CoV-2 infection may correlate with the encephalopathies observed in some COVID-19 patients (Roe, 2020) and further highlights the need to unravel the cellular basis of the observed anosmia.

Golden Syrian hamsters have been successfully used as a model of SARS-CoV-1 infection (Roberts et al., 2005) and have also been recently shown to be a good model for SARS-CoV-2 as well (Sia et al., 2020). Indeed, the expression profile of ACE2, the entry receptor for SARS-CoV-2, is very similar in hamsters and humans (Luan et al., 2020). Anosmia in COVID-19 patients provided impetus to study the expression profile of ACE2 in the nasal cavity. Several studies show that ACE2 is present in the OE but expressed in sustentacular cells, the supporting cells surrounding the olfactory neurons, rather than in the olfactory neurons themselves (Bilinska et al., 2020; Fodoulian et al., 2020). However, a recent study suggested that SARS-CoV-2 could infect olfactory sensory neurons in hamsters (Sia et al., 2020).

In the present work, we focused on the pathological impact of two different SARS-CoV-2 strains on the nasal cavity and the central nervous system of the golden Syrian hamster.

## 2. Material and methods

### 2.1 Isolates of SARS-CoV-2

Viral strains were isolated from nasopharyngeal swabs obtained from patients suffering from respiratory infection, suspected of COVID-19 and submitted to molecular diagnosis. Nasopharyngeal flocked swabs were suspended in UTM media (Copan, Italy), and kept at 4°C for less than 48 hours. A volume of 400 μl was homogenized and microfiltered (0.5 μm) previous to inoculation on Vero cells CCL-81 (passage 32, from ATCC, USA) grown at 80% confluence in a BSL3 virology laboratory (Virology Unit, CHU de Caen, France). Supernatants were harvested at day 3 after inoculation and immediately used in subsequent passage one (P1) of the virus following the same protocol. P1 was used for stock production and each stock was aliquoted and conserved at −80°C before titration and genomic quantification. Two strains were used in the model : UCN1 and UCN19, isolated at one week interval during the course of the active epidemic in Normandy, France (end of march).

### 2.2 Animals

Animal experiments were carried out in the animal-biosafety level 3 (A-BSL3) facility of the Animal Experimental Platform (ANSES – laboratoire de Lyon). The protocol for animal use was approved by the French Ministère de la Recherche et de l’Enseignement Supérieur number (Apafis n°24818-2020032710416319). Twelve female golden Syrian hamsters (eight weeks old) were purchased from Janvier’s breeding Centre (Le Genest, St Isle, France). Infection was done by nasal instillation (20μl in each nostril) of anaesthetised animals (Vetflurane). Five animals were infected with 5.4 10^3^ plaque forming units (pfu) of SARS-CoV2 strain UCN1, five animals were infected with 1.8 10^3^ pfu of SARS-CoV2 strain UCN19 and two mock-infected animals received only Dulbecco’s minimal essential medium (DMEM). SARS-CoV2-infected hamsters were euthanized after general anaesthesia (Vetflurane) at 2, 4, 7, 10 or 14 days after infection. At post-mortem, lungs were harvested, weighed and homogenized in 1 ml of DMEM with stainless steel beads (QIAGEN) for 30s at 30 Hz using TissueLyserII (QIAGEN). Lung homogenates were clarified by centrifugation (2,000 × g at 4°C for 10 min), aliquoted and stored at −80°C until RNA extraction. RNA extraction was done with QIAamp Viral RNA minikit (Qiagen) following manufacturer’s instructions. Real time quantitative PCR was performed on the E gene as described recently (Corman et al., 2020) on a Lightcycler LC480 (Roche).

### 2.3 Immunohistochemistry, histological stainings and quantifications

The immunohistochemistry analysis of the olfactory mucosa tissue sections was performed as described previously in mice (Bryche et al., 2019a). Briefly, the whole animal head was fixed for 3 days at room temperature in 4% paraformaldehyde PBS, then decalcified for one week (10% EDTA – pH 7.3 at 4°C). The nasal septum and endoturbinates were removed as a block and post-fixed overnight at 4°C in 4% paraformaldehyde PBS and further decalcified for 3 days. The same procedure was applied with cortex and brainstem but without decalcification. Blocks and tissues were cryoprotected with sucrose (30%). Cryo-sectioning (14 μm) was performed on median transversal sections of the nasal cavity, perpendicular to the hard palate, in order to highlight vomeronasal organ, olfactory mucosa, Steno’s gland and olfactory bulb. The brain was cryo-sectioned (14 μm) in the coronal plane. Sections were stored at −80°C until use. Non-specific staining was blocked by incubation in 2% bovine serum albumin and 0.3% Triton X-100. The sections were then incubated overnight with primary antibodies directed against SARS Nucleocapsid Protein (1:200; rabbit polyclonal, NB100-56576, NOVUS, France); olfactory marker protein - OMP (1:500; goat polyclonal; 544-10001, WAKO, USA), cytokeratin-18 - CK18 (1:50; mouse polyclonal; MAB3234 – RGE53, Sigma-Aldrich, France), ionized calcium-binding adapter molecule 1 - Iba1 (1: 1000; goat polyclonal; ab178846, Abcam, France); G alpha s/olf - G_olf_ (1:500; rabbit polyclonal; Santa Cruz Biotechnology (sc383), Dallas, TX, USA). Fluorescence staining was performed using goat anti-rabbit Alexa-Fluor-488 (1:1000; Molecular Probes – A32731; Invitrogen, Cergy Pontoise, France), donkey anti-goat Alexa-Fluor-555 (1:1000; Molecular Probes A32816; Invitrogen, Cergy Pontoise, France) and donkey anti-mouse Alexa-Fluor 555 (1:500; Molecular Probes – A32773; Invitrogen, Cergy Pontoise, France) secondary antibodies. Images were taken at x100 magnification using an Olympus IX71 inverted microscope equipped with an Orca ER Hamamatsu cooled CCD camera (Hamamatsu Photonics France, Massy, France) or with a Zeiss LSM 700 confocal 187 microscope (MIMA2 Platform, INRAe). Images were quantified using ImageJ (Rasband, W.S., ImageJ, U. S. National Institutes of Health, Bethesda, Maryland, USA, http://imagej.nih.gov/ij/, 1997–2012) to threshold specific staining.

For histology, some sections were rehydrated then stained in hematoxylin (H-3404 Vector Laboratories) for 30s and washed with water. Slides were then transferred in an acid-alcohol solution (0.5% HCl in 70% Ethanol) for 10s and washed in water, then immersed in eosin (Sigma HT110316) for 15s and washed again in water. Finally, slides were gradually dehydrated and mounted in Eukitt (Sigma 25608–33-7). Nissl staining was performed on some sections following conventional histological procedures, including immersion in 0.1% cresyl violet solution for 10 minutes. Images were taken at ×100 magnification using a DMBR Leica microscope equipped with an Olympus DP-50 CCD camera using Cell^F^ software (Olympus Soft Imaging Solutions GmbH, OSIS, Munster, Germany).

We measured the OE thickness, the OSN cilia quality (based on Golf staining) and the immune cell infiltration of the olfactory mucosa (based on the Iba1 staining) on 6 images per animal taken from 3 different slides spread along the nasal cavity. For the OE thickness, 10 measurements along the OE septum were performed for each image, as previously described (Francois et al., 2016). The percentage of apical OE with a preserved G_olf_ staining and the percentage of Iba1+ cells infiltration in the OE were measured using ImageJ to threshold specific staining. All values were averaged per animal.

We also globally estimated the impact of SARS-CoV-2 on the OE on these parameters. By comparing mock-infected animals with SARS-CoV-2 infected animals, we evaluated: ^1^the integrity of the structure of the OE using HE staining; ^2^the preservation of the OSN cilia using G_olf_ signal in the apical part of the OE; ^3^the presence and shape of Iba1+ cells in the OE and in the underneath *lamina propria.* A ramified Iba1+ cells is considered representative of a resting state while an amoeboid shape is linked to an activated state in the OE (Mori et al., 2002). For each parameter, we used a scale ranging from 0 to 10 scored on three sections of the olfactory mucosa spread along the nasal cavity per animal. This evaluation was performed by three experienced investigators blinded to the animal treatment. Scores were averaged per animal between investigators.

## 3. Results

### 3.1 SARS-CoV-2 induces massive damage in the olfactory epithelium

We used a cohort of 12 hamsters to examine the impact of two different isolates of SARS-CoV-2 (UCN1 and UCN19) on the nasal cavity from 2 to 14 day post infection (DPI) through nasal instillation. While the olfactory epithelium was well preserved in control animals, we observed massive damage of the olfactory epithelium in SARS-CoV-2-infected animals as early as 2 DPI (**Fig. 1A**). The nasal cavity lumen was filled with cellular aggregates and most of the olfactory epithelium was strongly disorganized. In the most affected OE zones, axon bundles from the olfactory sensory neurons under the olfactory epithelium were almost in direct contact with the environment (**Fig. 1B**). As an objective measure of olfactory mucosa damage, we first measured the septal olfactory epithelium thickness (**Fig. 1C**) and we then scored the overall lesions of the OE (**Fig. 1D**). Most of the OE had disappeared at 4 DPI, after which point it started to recover but had not reached pre-infection thickness at 14 DPI when our measurements stopped. The results were similar for both viruses.

**Figure 1:**
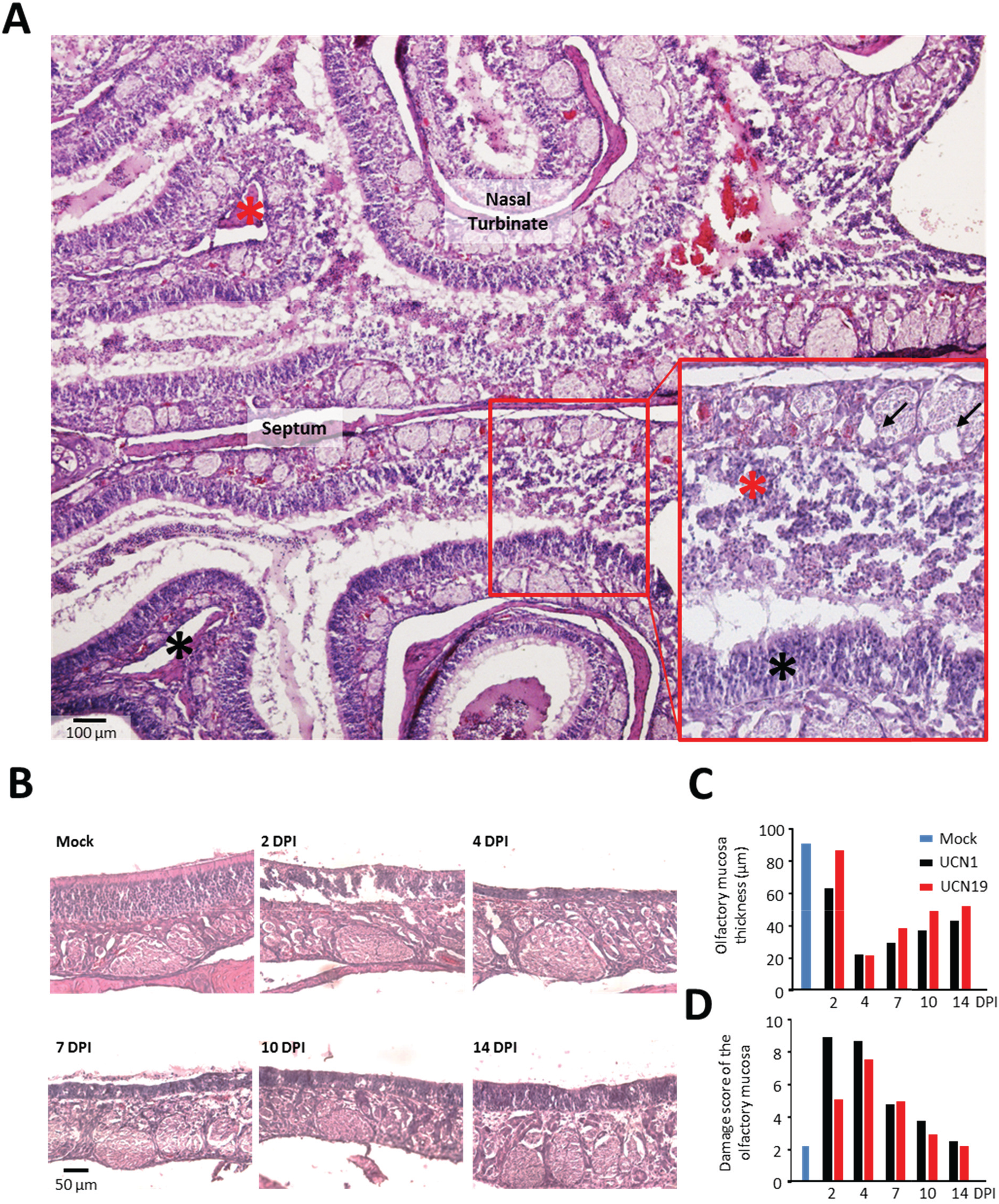
SARS-CoV-2 nasal instillation induces massive damage of the olfactory epithelium 2 DPI which has partially h aled 14 DPI. **A** Global view of the olfactory nasal turbinates surrounding the septum (2 DPI, Virus UCN19). Some parts of the olfactory epithelium are mildly damaged (black asterisk) while other parts are totally destroyed and released into the lumen of the nasal cavity (red asterisk), leaving the lamina propria in direct contact with the environment along with axon bundles (arrows). **B** Evolution of the dorsal septal region of the nasal cavity from mock infected hamsters to 14 DPI in UCN19 virus-infected hamsters. **C** Thickness of the olfactory epithelium in the same region. **D** Damage score of the olfactory epithelium for both viruses. Each histogram represents a different animal except for mock (n=2).

### 3.2 SARS-CoV-2 infects mainly sustentacular cells and not olfactory sensory neurons in the OE

We then examined the cellular localisation of SARS-CoV-2 in the nasal cavity by immunohistochemistry, using an antibody raised against the nucleocapsid protein of the virus. We observed that a large proportion of the OE was infected at 2 DPI, which decreased at 4 DPI and subsequently disappeared (**Fig. 2A and Supp. Fig. 1A, B**). The virus was also present in the vomeronasal organ (**Supp. Fig. 2A**) and in the Steno’s gland (**Supp. Fig. 2B**) at 2 DPI and 4 DPI. We also examined various areas of the brain. We could not find any presence of the virus in the olfactory bulbs (**Supp. Fig. 3A**), in the olfactory cortex (piriform cortex and olfactory tubercle), hypothalamus, hippocampus, and in the brainstem where we mainly focused on the respiratory control centres: the ventral respiratory column and the nucleus of the solitary tract (**Supp. Fig. 3B**). The virus presence in the nasal cavity followed a kinetic similar to its presence in the lungs (**Supp. Fig. 1C**).

**Figure 2:**
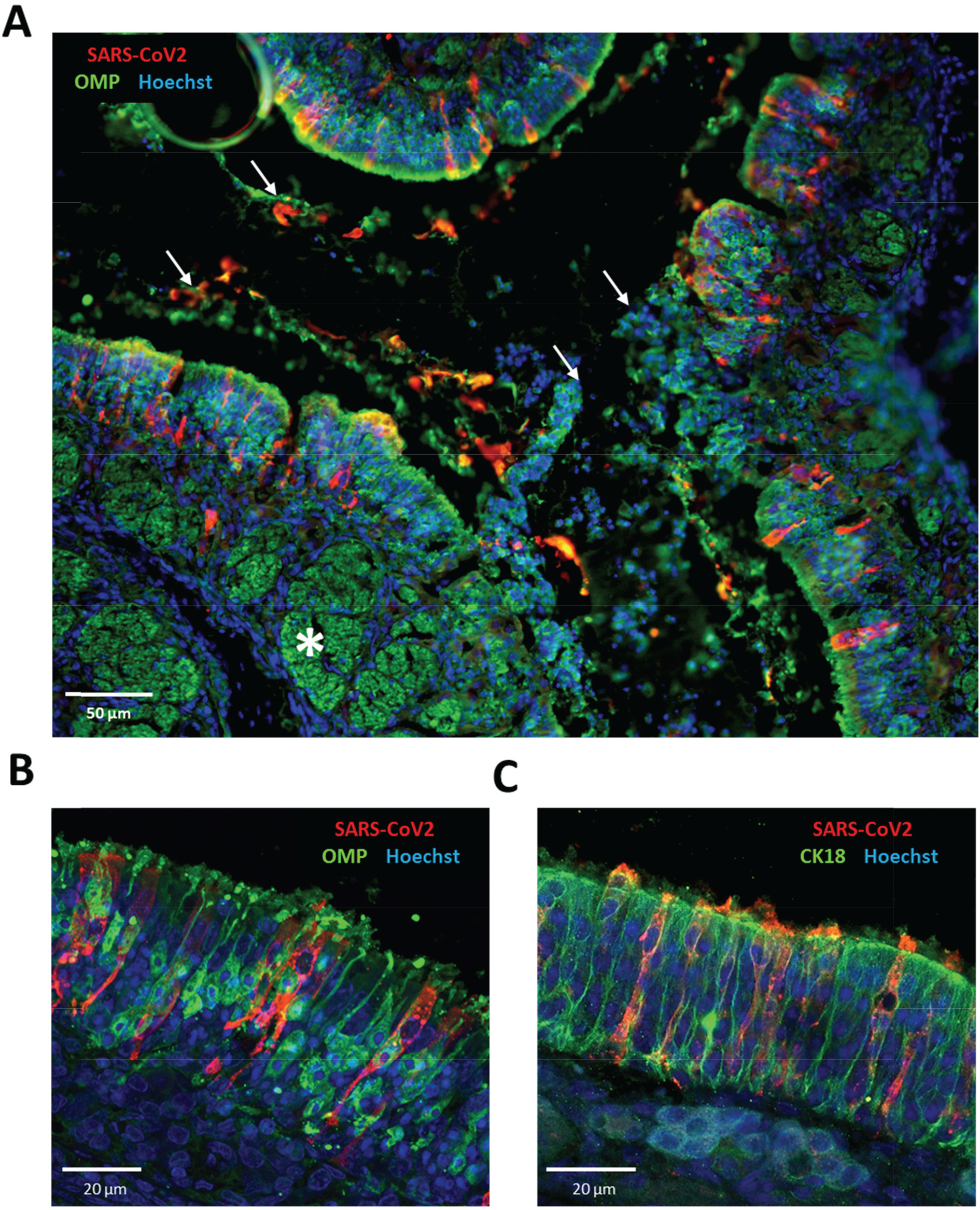
Localization of SARS-CoV-2 in the nasal cavity. **A** Representative image of double staining of the olfactory mucosa against SARS-CoV-2 and a marker of olfactory sensory neurons (OMP, 4 DPI, UCN19). The OE is mostly disorganized but axon bundles expressing OMP are well preserved (white asterisk). The nasal cavity lumen contains many cellular aggregates revealed by the nuclear Hoechst staining along with the presence of SARS-CoV-2 antigens and OMP (white arrows). The proximity to the damaged OE suggests that these cellular aggregates could originate from the disassociation of the OE from the lamina propria. **B** Same double staining in a relatively preserved area of the OE from the same animal showing an absence of overlap between OMP and SARS-CoV-2 staining. **C** Representative image of double staining of the olfactory mucosa against SARS-CoV-2 and a marker of sustentacular cells (CK18, 4 DPI, UCN19) which colocalize.

In order to identify which cells were infected with SARS-CoV-2 in the OE, we performed double staining of the olfactory mucosa with OMP and CK18, specific markers of mature OSNs and sustentacular cells respectively. While most of the cells infected with SARS-CoV2 expressed CK18 as well, we were not able to find any viral antigen in OMP expressing cells. These results show that SARS-CoV-2 infects sustentacular cells but not OSNs (**Fig. 2**). Interestingly, we observed staining of OMP, CK18, SARS-CoV-2 as well as Hoechst in the lumen of the nasal cavity indicating that part of the OE containing infected and non-infected cells was desquamated in response to SARS-CoV-2 infection.

### 3.3 OSNs lost most of their cilia as early as 2 DPI with SARS-CoV-2

In order to understand the extent of the damage of the OSNs following SARS-CoV-2 infection, we examined the preservation of the OSN cilia layer, which contains all the transduction complex allowing the detection of odours (Dibattista et al., 2017). We performed immunohistochemistry against G_olf_ (**Fig. 3A, B**), a specific G protein from this complex (Jones and Reed, 1989). We quantified the percentage of septal olfactory epithelium containing G_olf_ staining in the apical part of the OE (**Fig. 3C**) and scored globally the quantity of G_olf_ staining in the OE (**Fig. 3D**). While the G_olf_ signal was completely preserved in control animals, it quickly diminished in SARS-CoV-2-infected animals without any clear improvement from 2 DPI to 10 DPI. We observed a small recovery of the G_olf_ signal only in the septal part of the OE 14 DPI with the UCN19 virus.

**Figure 3:**
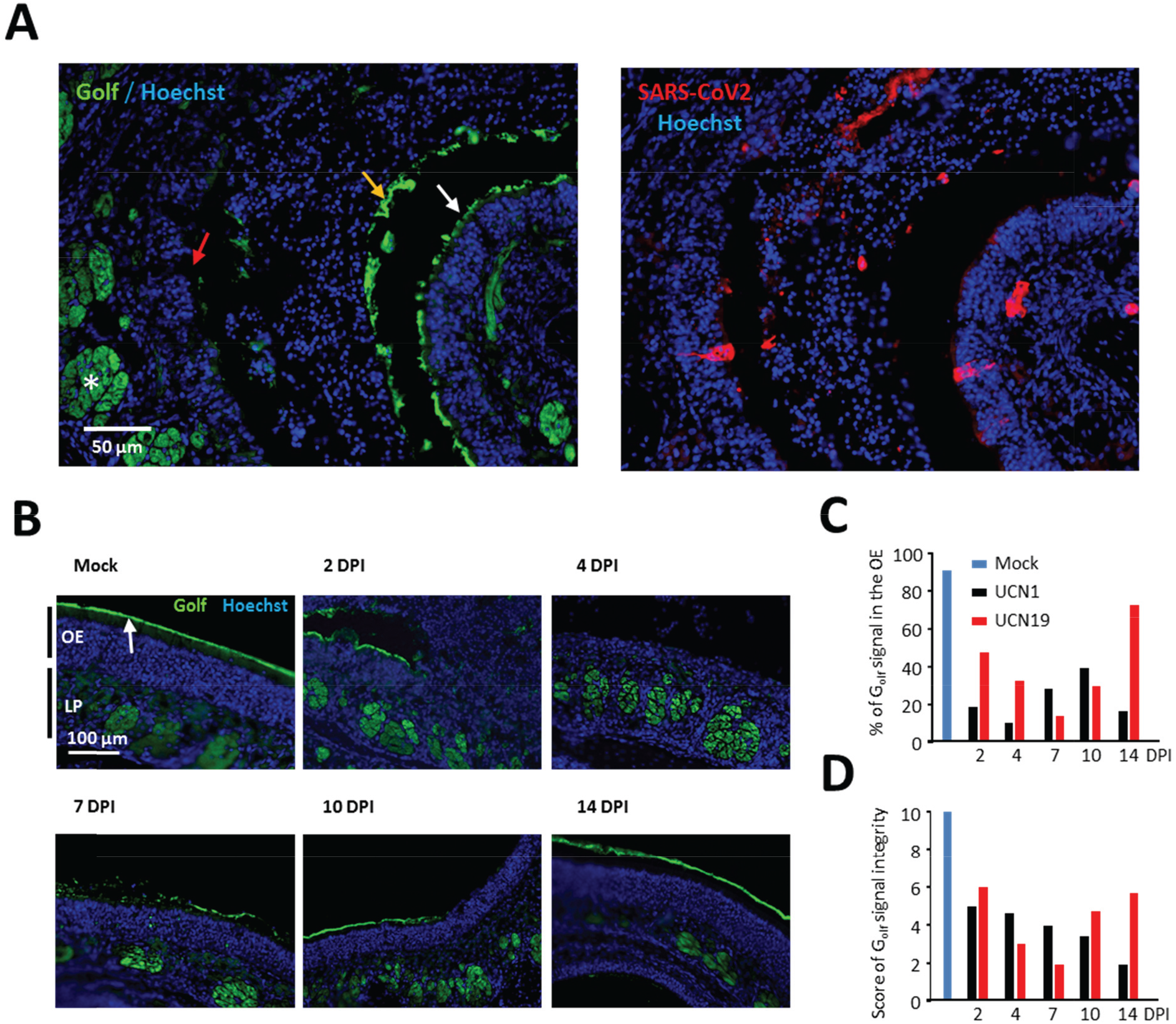
OSNs cilia are severely impaired following SARS-CoV-2 infection in the nasal cavity. **A** Representative images of immunostaining against G_olf_ (left) and SARS-CoV-2 (right) on two successive slides of the olfactory mucosa (4DPI, UCN19). Some parts of the OE still hold some of G_olf_ staining (white arrow), but most of it was eliminated in the cellular aggregates present in the nasal cavity lumen (orange arrow). Other OE apical regions were devoid of G_olf_ signal (red arrow) while retaining the signal in the axon bundles (white asterisk) **B** Representative images of G_olf_ staining from mock treated animals and 2 to 14 DPI in UCN19 virus-infected hamsters. Olfactory epithelium (OE) and lamina propria (LP) are indi ated on the left of the first image. The cilia layer stained with G_olf_ signal is indicated by a white arrow. **C** Percentage of apical border of the septal OE stained with G_olf_. **D** Scores of G_olf_ presence in the apical part of the OE for both viruses. Each histogram represents a different animal except for mock (n=2).

### 3.4 Infiltration of immune cells in the olfactory epithelium following SARS-CoV-2 intranasal instillation

We finally examined the presence of immune cells in the olfactory mucosa using Iba1+ as a specific marker of monocyte/macrophage lineage (Bryche et al., 2019b); (**Fig. 4A**). In control animals, Iba1+ cells were mainly localized in the *lamina propria* and exhibited a ramified morphology with many processes (**Fig. 4B**), which is typical of a resting phenotype (Mori et al., 2002). We measured the presence of Iba1+ cell (**Fig. 4C**) and scored the presence and shape of Iba1+ cells in the OE and the *lamina propria* separately (**Fig. 4D**). While Iba1+ cells were mostly absent from the OE and moderately present in the lamina propria in control animals, their presence was drastically increased up to 4 DPI in infected animals, both in the OE and the *lamina propria* (**Fig. 4A, B**). Furthermore, Iba1+ cells were mostly of amoeboid shape which is typical of an activated state at 2 to 7 DPI (**Fig. 4B**). We also observed Iba1+ cells in the lumen of the nasal cavity in the cellular aggregates containing SARS-CoV-2 (**Fig. 4A**). Immune cell infiltration gradually diminished. This reduction was first observed at 10 DPI in the OE but was not seen in the *lamina propria* before 14 DPI. At 14 DPI, it reached the mock level both in the OE and the *lamina propria* for both virus strains (**Fig. 4C, D**).

**Figure 4:**
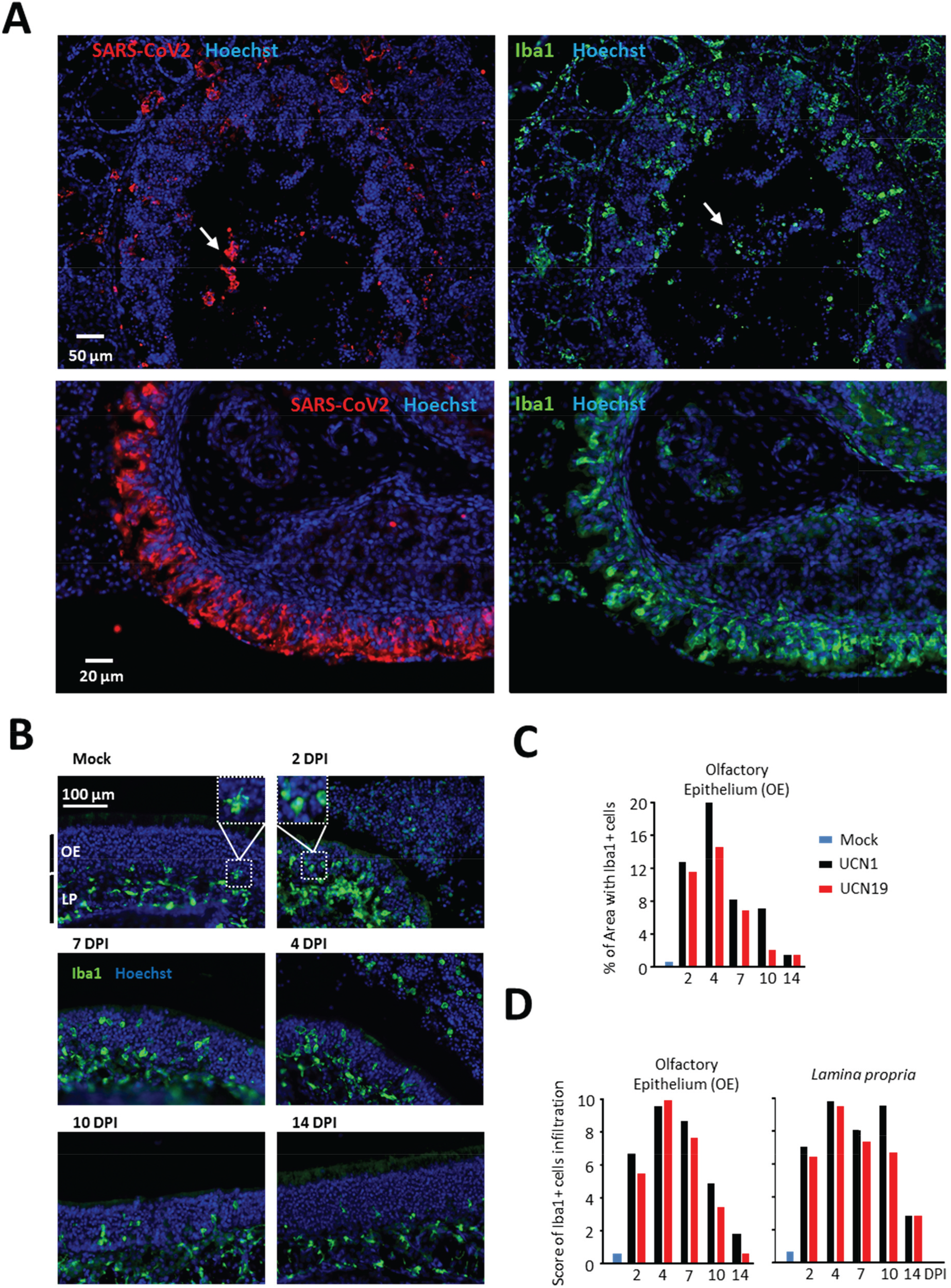
Infiltration of immune cells in the olfactory mucosa following SARS-CoV-2 infection. **A** Representative images from successive slides of immunostaining against Iba1 and SARS-CoV-2 (2 DPI, UCN19). Iba1+ cells are massively present in both the olfactory epithelium and the underlying lamina propria. Both Iba1+ cells and SARS-CoV-2 are also present in the cellular aggregates in the lumen of the nasal cavity near the disorganized olfactory epithelium (upper panels, white arrows). We also observed a similar localization of infected cells and immune cells in relatively preserved part of the olfactory epithelium (lower panel). **B** Representative images of Iba1 staining from mock treated animals and 2 to 14 DPI in infected animals (UCN19). Olfactory epithelium (OE) and lamina propria (LP) are indicated on the left of the first image. The typical phenotypes of a resting Iba1+ cell presenting ramified shape (mock) and of an active phenotype of amoeboid shape are presented in the upper panel of the image. **C** Percentage of Iba1+ cells area in the olfactory epithelium. **D** Scores of Iba1+ cells presence and shape in the olfactory epithelium (OE) and lamina propria (LP). Each histogram represents a different animal except for mock (n=2).

## 4. Discussion

In the present study, we focused on the impact of SARS-CoV-2 in the nasal cavity and central nervous system using the golden Syrian hamster as a model. We chose to work with 2 different isolates of SARS-CoV-2, with the potential to obtain slightly different strain behaviour. We explored the kinetic of the impact of these two different isolates. Due to the difficulty of designing experiments during the COVID-19 pandemic, we examined only one animal per virus strain and per time point. Overall, our results are globally consistent for both viruses and the kinetics of the different measured parameters were always following consistent evolution from 2 DPI to 14 DPI. While these results require confirmation with larger animal groups, they provide a qualitative overview of the impact of SARS-CoV-2 on the nasal cavity which could explain the anosmia observed in COVID-19 patients. Indeed, according to our results, sustentacular cells are rapidly infected by SARS-CoV-2 and this infection is associated with a massive recruitment of immune cells in the OE and lamina propria, which could induce a rapid disorganization of the OE structure by causing immunopathology. This hypothesis is consistent with elevated level of TNFα observed in OE samples from COVID-19 suffering patients (Torabi et al., 2020). This disorganization is followed by a massive destruction of the OE which is released in the lumen of the nasal cavity, taking away most of the cilia layer of the OSNs. In the absence of these cilia, the olfactory transduction is not functional and thus the olfactory capacity of the animals should be impaired until the OE regenerates. At 14 DPI, the thickness of the OE is already recovering to approximatively 50% of the mock condition and some functional cilia reappear. Such kinetics is consistent with the observed recovery of anosmia in COVID-19 patients. Indeed, recent studies highlight a relatively fast recovery (~15 days) (Dell'Era et al., 2020; Meini et al., 2020), compatible with the observed partial recovery of the OE in hamsters 14 DPI.

We identify sustentacular cells as the main target cells of SARS-CoV-2 in the OE. This observation is consistent with the expression pattern of ACE2 mainly present in these cells along with the facilitating protease TMPRSS2 (Bilinska et al., 2020; Fodoulian et al., 2020). Two recent reports indicate that olfactory neurons in hamster (Sia et al., 2020) and respiratory cells in ferret (Ryan et al., 2020) may be the target of SARS-CoV-2 but these studies did not focus on the nasal cavity and they did not use double staining to clearly identify the infected cells in the OE. We observed that SARS-CoV-2 can infect other non-neuronal cells in the nasal cavity, notably in the epithelium covering the lumen of Steno’s gland which is however poorly described in the literature (Bryche et al., 2019b). Further studies are required to clearly identify the other non-neuronal cells infected by SARS-CoV-2 in the nasal cavity which may facilitate the systemic infection of other cell types and tissues lower in the respiratory airways.

We did not observe any presence of the virus in the brain, notably in the olfactory bulbs where OSNs projects and in the piriform cortex where the olfactory signal is integrated. We also focused on the hypothalamus which contains ACE2 expressing neurons (Mukerjee et al., 2019) as well as on the respiratory centres of the brainstem as the latter are suspected to be infected in COVID-19 patients suffering from heavy respiratory disorders (Gandhi et al., 2020). Our lack of virus detection in the central nervous system may be due to the low number of animal examined but, nevertheless, we can rule out a systematic and important infection of the brain following SARS-CoV-2 infection in hamster. This is consistent with a recent review of the literature which failed to indicate any central nervous manifestation of SARS-CoV-2 presence in human central nervous system (Romoli et al., 2020), however the neurotropic ability of SARS-CoV-2 remains controversial (Wang et al., 2020). Interestingly, SARS-CoV-1 has been shown to be neurotropic only using ACE2 humanized mice. The OSNs of these mice must also express ACE2 as it is under the control of K18 ubiquitous promoter (Netland et al., 2008). SARS-CoV-1 may thus infect OSNs expressing ectopic ACE2 and the observation of presence of the virus in the brain may not be relevant for a more physiological model. In our model, we observed a fast desquamation of the OE following SARS-CoV-2 infection of sustentacular cells. Further studies are required to decipher whether it could be an anti-viral strategy to limit access of the virus to the brain through the olfactory pathway and to which extent the recruitment of immune cells contributes to this process through immunopathological mechanisms.

Globally, we present here the first data focusing on the impact of the SARS-CoV-2 in the nasal cavity. Our results could explain the high prevalence of anosmia observed in COVID-19 patients and will need to be confirmed by analysis of human OE.

## Abbreviations

OSN: Olfactory sensory neuron
OE: Olfactory epithelium

## Acknowledgement

We would like to thank Pr. Astrid Vabret (head of the Laboratoire de Virology, CHU de CAEN) for BSL3 facilities access and isolates production, Estelle Leperchois (CHU de Caen) for cell production, Latifa Lakhdar (head of the Plateforme d’expérimentation animale, Anses - Laboratoire de Lyon), Bruno Da Costa and Christophe Chevalier for their help with handling samples, Bernard Delmas and all members of VIM for helpful discussions, Pierre Adenot for exceptional access to the confocal during the pandemic (MIMA2 platform, https://doi.org/10.15454/1.5572348210007727E12) and Birte Nielsen for her help improving the manuscript. This work has been submitted to Brain, Behavior, and Immunity Journal the 16^th^ of June 2020.

---

**Supplementary Figure 1:**
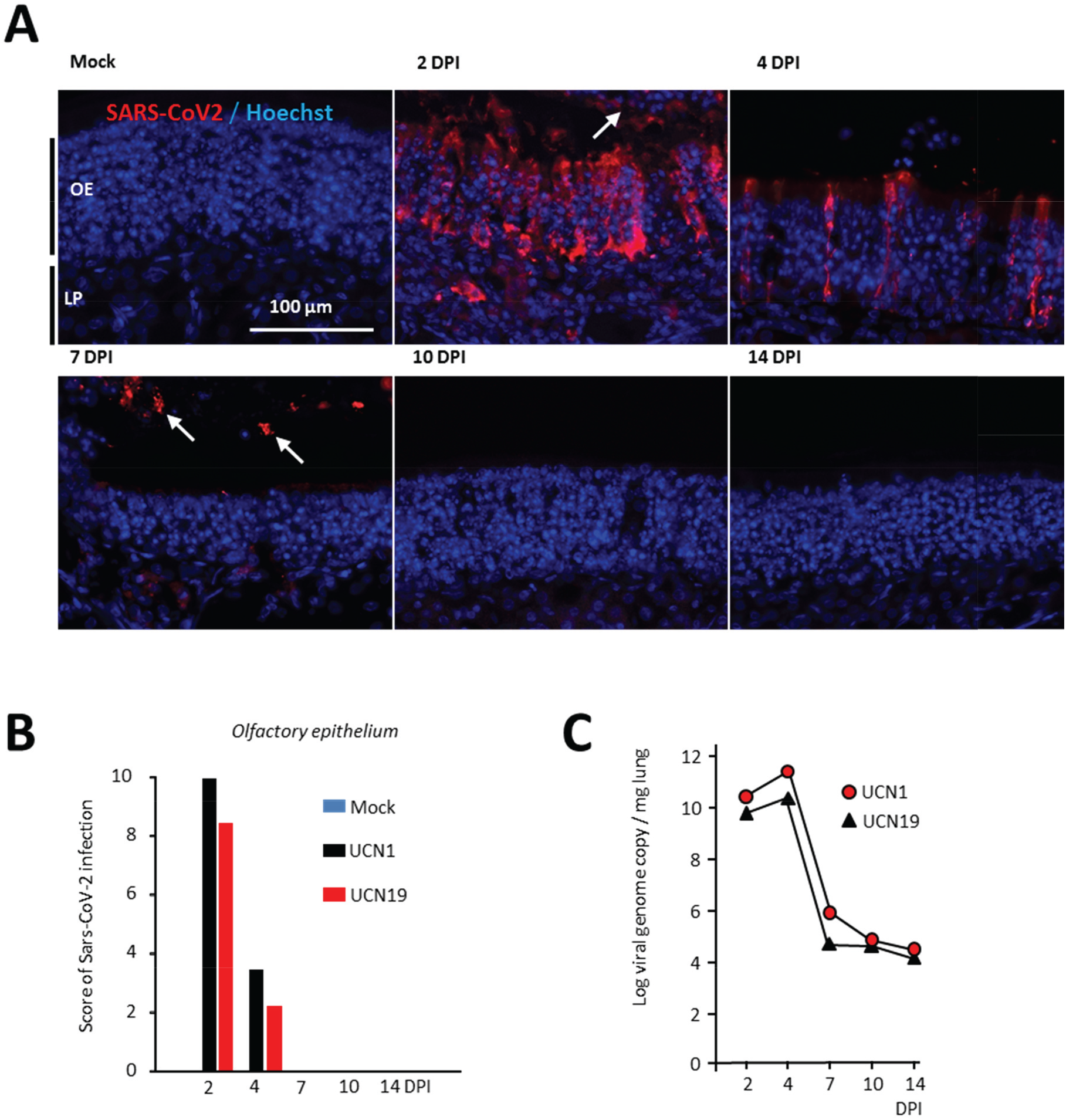
**(A)** Representative images of the presence of SARS-CoV-2 in the olfactory epithelium from Mock to 14 DPI (white arrows indicate the presence of the virus in the lumen of the nasal cavity) with (**B**) associated score of presence in the olfactory epithelium. Olfactory epithelium (OE) and *lamina propria* (LP) are indicated on the left of the first image. (**C**) Presence of the virus in the lungs.

**Supplementary Figure 2:**
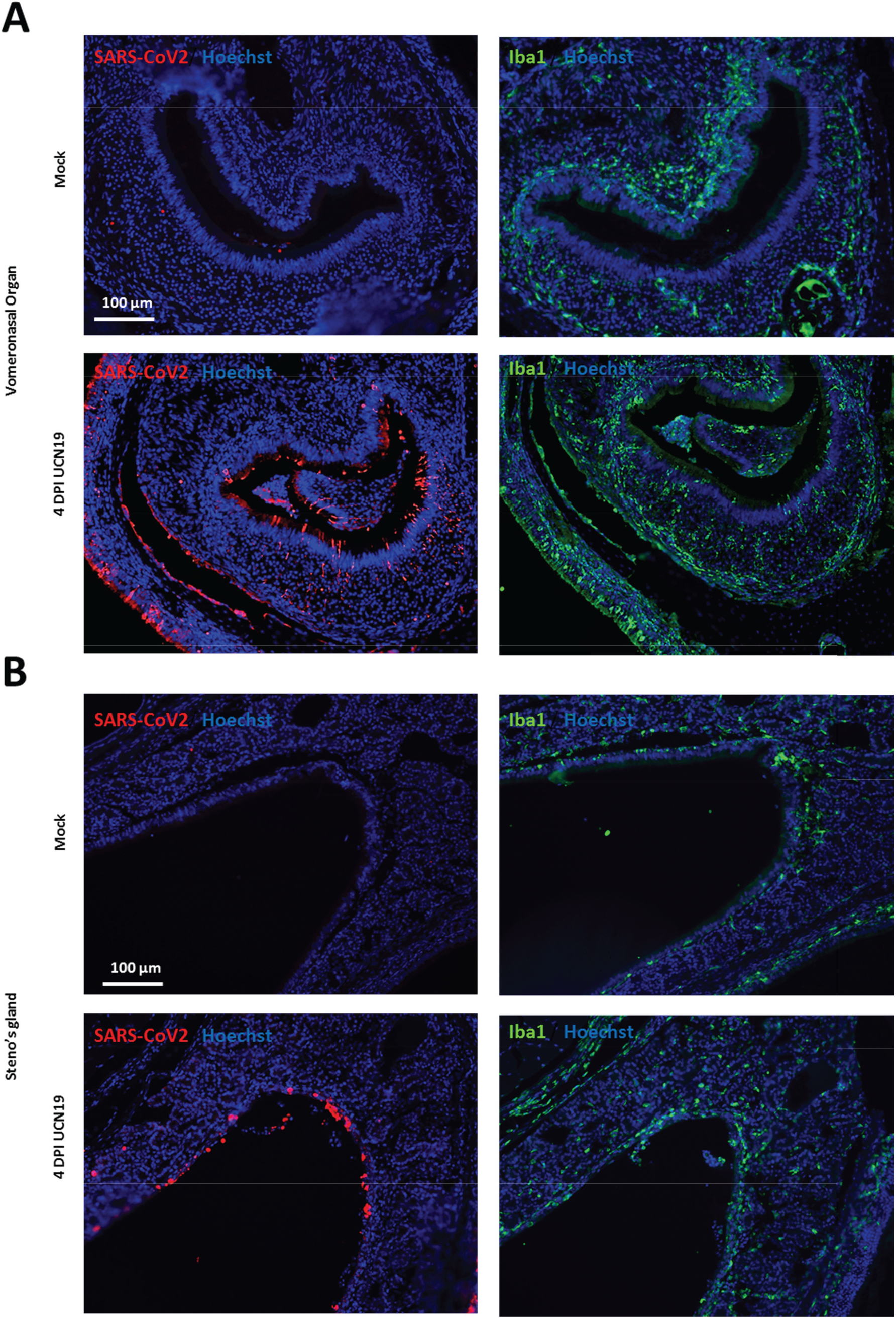
Presence of SARS-CoV-2 and immune cells outside of the olfactory epithelium in the nasal cavity. (**A**) In the vomeronasal organ and (**B**) in the Steno's gland.

**Supplementary Figure 3:**
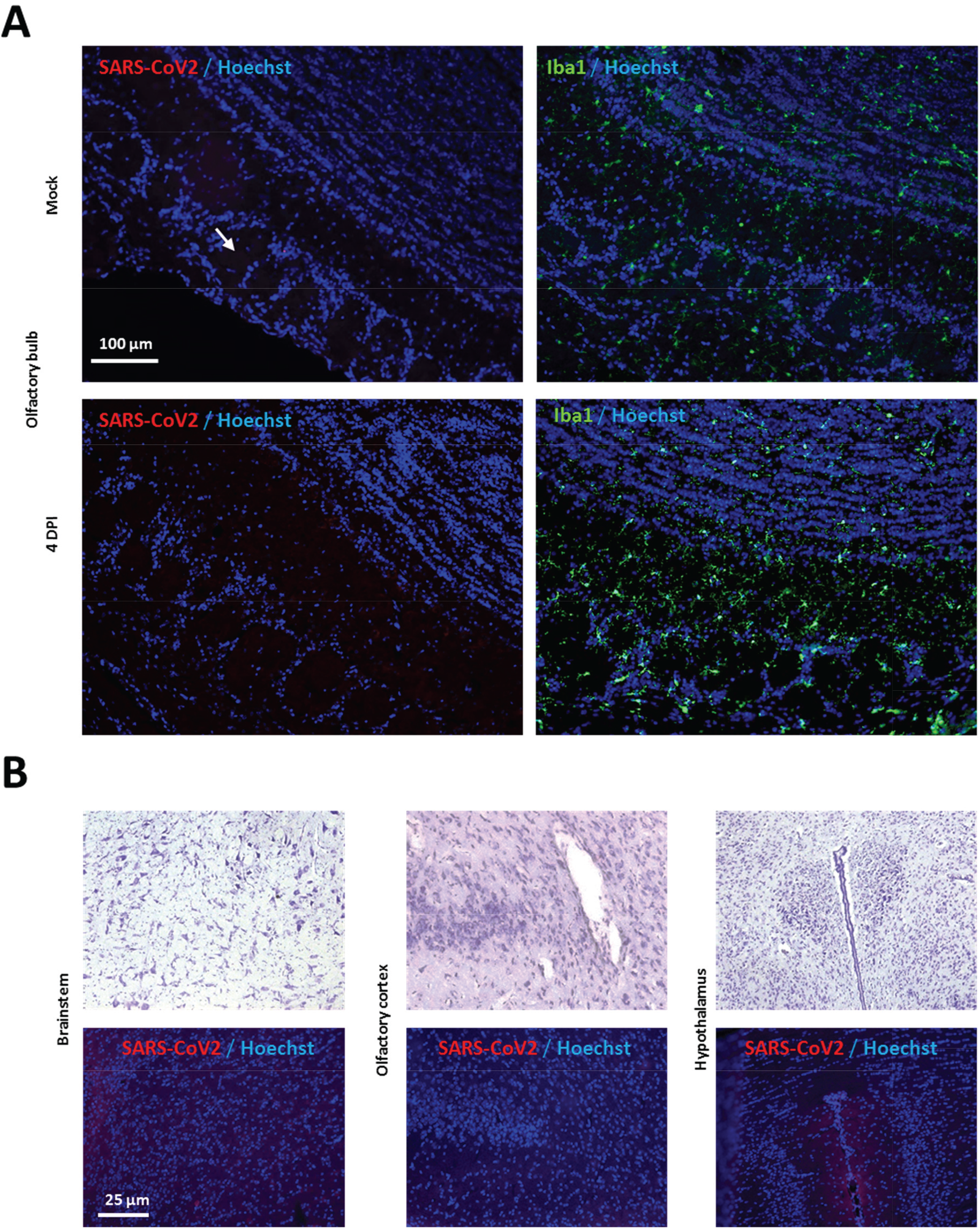
Presence of SARS-CoV-2 and immune cells (Iba1+) in the centra nervous system (J4, ICN19). (**A**) In the olfactory bulb (white arrow indicates a glomerulu**s**) and (**B**) in different areas of the central nervous system (Upper image is the Nissl stain of the corresponding area on successive slide): Brainstem (focus on the ventral respiratory column and the nucleus of the solitary tract); Olfactory cortex (piriform cortex and olfac ory tubercle), hypothalamus (paraventricular nucleus).

